# Optineurin mediates delivery of membrane from transferrin receptor-positive endosomes to autophagosomes

**DOI:** 10.1101/2022.01.12.476135

**Authors:** Megha Bansal, Kapil Sirohi, Shivranjani C Moharir, Ghanshyam Swarup

## Abstract

Autophagy is a conserved quality control mechanism that removes damaged proteins, organelles and invading bacteria through lysosome-mediated degradation. During autophagy several organelles including endoplasmic reticulum, mitochondria, plasma membrane and endosomes contribute membrane for autophagosome formation. However, the mechanisms and proteins involved in membrane delivery to autophagosomes are not clear. Optineurin (OPTN), a cytoplasmic adaptor protein, is involved in promoting maturation of phagophores into autophagosomes; it is also involved in regulating endocytic trafficking and recycling of transferrin receptor (TFRC). Here, we have examined the role of optineurin in the delivery of membrane from TFRC-positive endosomes to autophagosomes. Only a small fraction of autophagosomes was positive for TFRC, indicating that TFRC-positive endosomes could contribute membrane to a subset of autophagosomes. The percentage of TFRC-positive autophagosomes was reduced in Optineurin knockout mouse embryonic fibroblasts (*Optn^−/−^* MEFs) in comparison with normal MEFs. Upon over-expression of optineurin, the percentage of TFRC-positive autophagosomes was increased in *Optn^−/−^* MEFs. Unlike wild-type optineurin, a disease-associated mutant, E478G, defective in ubiquitin binding, was not able to enhance formation of TFRC-positive autophagosomes in *Optn^−/−^* MEFs. TFRC degradation mediated by autophagy was decreased in optineurin deficient cells. Our results suggest that optineurin mediates delivery of TFRC and perhaps associated membrane from TFRC-positive endosomes to autophagosomes, and this may contribute to autophagosome formation.

## Introduction

Optineurin (OPTN) is a multifunctional cytoplasmic adaptor protein involved in signalling, membrane vesicle trafficking, autophagy, endoplasmic reticulum (ER) stress response and several other functions [1–8]. It is an autophagy receptor that is involved in mediating clearance of cytosolic *Salmonella*, damaged mitochondria and mutant proteins aggregates [7, 9–11]. Its functions are generally mediated by interaction with other proteins, often by linking two or more proteins. The autophagic function of OPTN depends on its interaction with the autophagosomal protein LC3 and ubiquitin through well-defined binding sites [7]. OPTN is also involved in mediating several membrane vesicle trafficking pathways between the Golgi and plasma membrane [12–18]. OPTN interacts with an activated form of Rab8 and mediates Rab8-dependent endocytic trafficking and recycling of transferrin receptor (TFRC) [13, 17]. It is required for trafficking of TFRC from early endosomes at the periphery of the cell to juxtanuclear region [15, 16]. It is localized in the Golgi, endosomes, autophagosomes, lysosomes and various vesicles involved in mediating transport [7, 13–18].

TFRC is a transmembrane protein involved in iron homeostasis. Binding of iron-bound transferrin to TFRC at the plasma membrane initiates the process of endocytosis leading to formation of TFRC-positive vesicles which move to early endosomes and then to recycling endosomes [19, 20]. Iron is released in the endosomes due to low pH. Most of the TFRC and transferrin is recycled back to plasma membrane and a small fraction is constitutively degraded in lysosomes through autophagy [21, 22]. A glaucoma-associated variant of optineurin, M98K, causes excessive autophagic degradation of TFRC, which leads to death of retinal cells [21, 22].

Autophagy is one of the mechanisms used by the cell to maintain homeostasis, that removes damaged proteins, organelles and invading bacteria through lysosomal degradation [23, 24]. Impaired autophagy is associated with several human disorders including cancer and neurodegenerative disorders [24]. Macroautophagy, here after referred to as autophagy, involves formation of phagophore (or isolation membrane), a cup-shaped double membrane structure [23, 24]. The phagophore grows and matures into a closed structure known as autophagosome, which fuses with lysosomes leading to formation of autolysosomes where degradation of the cargo takes place. Several autophagy related (Atg) proteins and signalling proteins regulate formation of phagophore and its maturation into autophagosome [23, 24, 26, 27]. The autophagosomal protein LC3 recruits the cargo, which needs to be degraded, to the autophagosomes. Membrane from several different intracellular organelles such as endoplasmic reticulum (ER), plasma membrane, Golgi, early and recycling endosomes, is delivered to autophagosomes during their biogenesis [28, 29]. However, the proteins and mechanisms involved in the delivery of membrane for autophagosome biogenesis are poorly understood. Whether membrane from a particular organelle contributes to a subset of autophagosomes or all the autophagosomes is not clear.

Since OPTN is involved in mediating various membrane vesicle trafficking pathways, it is a good candidate for facilitating delivery of membrane to autophagosomes that is required for their biogenesis. Previously, it has been shown that vesicles derived from TFRC containing Rab11-positive recycling endosomes are delivered to forming autophagosomes [28, 29]. However, the role of OPTN in the delivery of membrane from recycling endosome to autophagosome has not been investigated. OPTN is known to promote maturation of phagophore into autophagosome by enhancing LC3-II production [30]. Maturation of phagophore into autophagosome also requires delivery of membrane from various organelles. Here, we have investigated the role of OPTN in the delivery of membrane from TFRC-positive endosomes to autophagosomes. Our results show that formation of transferrin receptor-positive autophagosomes is reduced in OPTN deficient cells compared to normal cells, whereas overexpressed OPTN enhances formation of transferrin receptor-positive autophagosomes. Autophagy mediated TFRC degradation was also reduced in optineurin deficient cells. A disease-associated mutant of OPTN, E478G, that is defective in autophagy, is also defective in enhancing the formation of TFRC-positive autophagosomes.

## Results

### Overexpressed OPTN enhances formation of TFRC-positive autophagosomes

During autophagy, TFRC-positive recycling endosomes contribute membrane for the formation of autophagosomes [28, 29]. Since OPTN is involved in the regulation of endocytic trafficking and recycling of TFRC [15–17], we examined the role of OPTN in the delivery of membrane from TFRC-positive endosomes to autophagosomes. We hypothesised that if OPTN is involved in this process, then over-expression of OPTN would be expected to increase the formation of TFRC-positive autophagosomes. We expressed GFP-LC3 along with either HA-tagged OPTN, or empty vector in OPTN knockout mouse embryonic fibroblasts (*Optn^−/−^* MEFs), and, after 24 hours of expression, fixed the cells and stained with antibodies against TFRC and HA. These cells were examined by confocal microscopy. We analysed the percentage of autophagosomes (GFP-LC3-positive dots) that were co-localising with endogenous TFRC (Fig. 1A, B). Presence of OPTN significantly increased the percentage of TFRC-positive autophagosomes from 12.57% to 34.29% in *Optn^−/−^* MEFs (Fig. 1B). The percentage of OPTN-positive autophagosomes was 43.22% and majority of the TFRC-positive autophagosomes were also positive for OPTN, as seen from the merged image (Fig. 1A).

**Figure 1.**
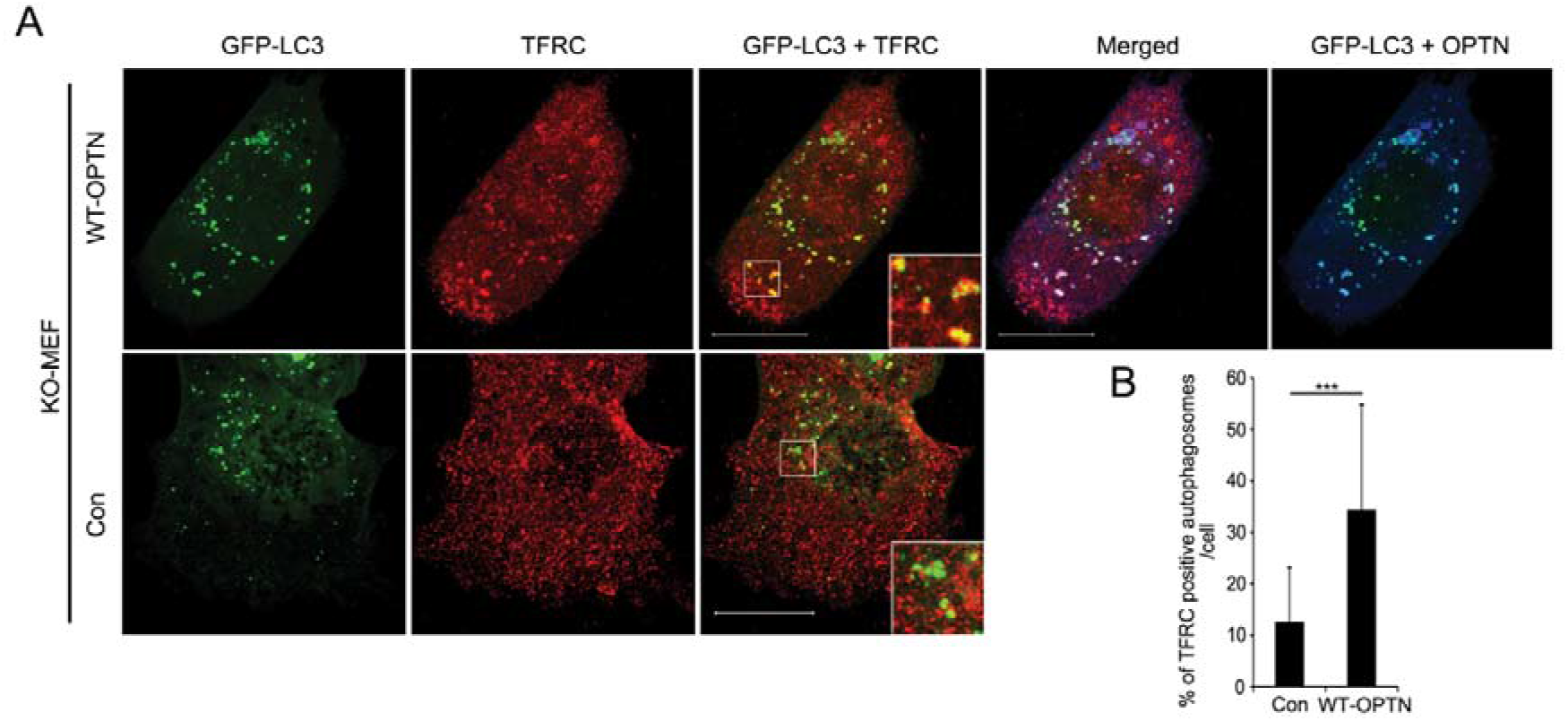
Overexpressed optineurin enhances formation of TFRC-positive autophagosomes. **(A)** Representative confocal images showing co-localization of endogenous TFRC with GFP-LC3B in *Optn^−/−^* (KO) MEFs transfected with GFP-LC3B along with either control plasmid (Con) or HA-WT-OPTN construct. TFRC was stained with anti-mouse Cy3 antibody (represented as red) while HA-OPTN was stained with HA antibody followed by anti-rabbit Alexa-633 antibody (represented as blue). Magnified area is shown as inset. Scale bar: 20μm. **(B)** Bar diagram showing percentage of GFP-LC3B-positive dots (autophagosomes) co-localizing with endogenous TFRC in *Optn^−/−^* MEFs transfected with GFP-LC3B along with either control plasmid (Con) or HA-WT-OPTN construct. n=42 cells. ***p<0.001.

### Formation of TFRC-positive autophagosomes is reduced in OPTN-deficient cells

To examine the role of endogenous OPTN in the formation of TFRC-positive autophagosomes, we determined the number and percentage of autophagosomes that were co-localizing with TFRC in *Optn^+/+^* and *Optn^−/−^* MEFs by immunostaining endogenous TFRC and endogenous LC3, after treating the cells with 25μM chloroquine for 12 hrs (Fig. 2 A, B). Treatment with chloroquine (which inhibits autophagy flux) was done to enhance staining of endogenous LC3. The percentage of LC3-positive dots (autophagosomes) that were also positive for TFRC was significantly less in *Optn^−/−^* MEFs compared to *Optn^+/+^* MEFs (Fig. 2 B). The average number of TFRC-positive autophagosomes was significantly higher (p<0.001) in *Optn^+/+^* cells (27.46/cell) compared to *Optn^−/−^* cells (4.63/cell). These results suggest that optineurin plays an important role in the formation of TFRC-positive autophagosomes.

**Figure 2.**
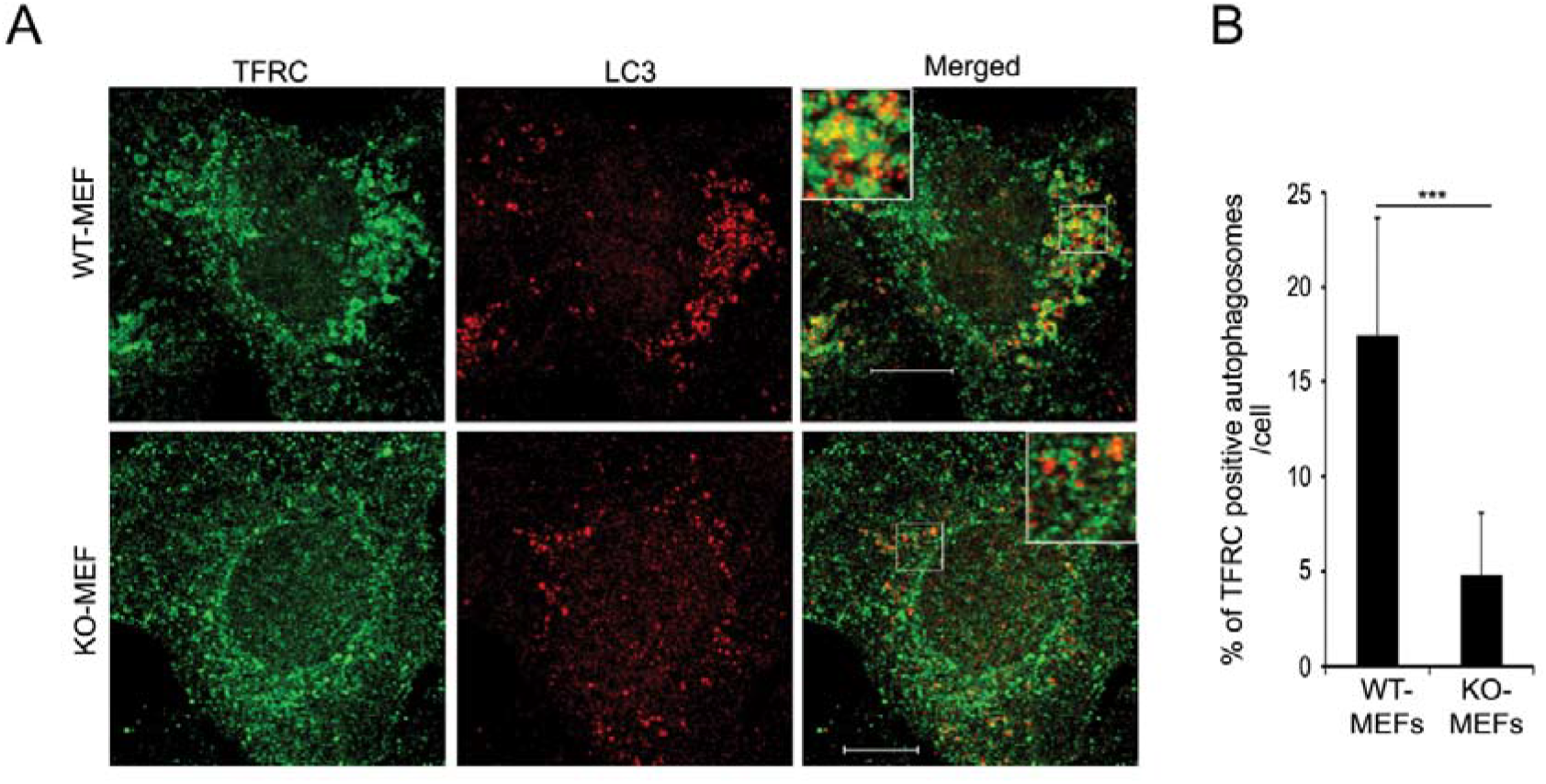
Formation of TFRC-positive autophagosomes is reduced in optineurin-deficient cells. **(A)** Representative confocal images showing co-localization of endogenous TFRC with endogenous LC3B in *Optn^+/+^* (WT) and *Optn^−/−^* (KO) MEFs treated with 25μM of chloroquine for 12 hrs. Magnified area is shown as inset. Scale bar: 10μm **(B)** Bar diagram showing percentage of autophagosomes per cell that are TFRC-positive in *Optn^−/−^* and *Optn^+/+^* MEFs treated with 25μM of chloroquine for 12 hours. n=11 cells. ***p<0.001.

### Autophagy mediated TFRC degradation is reduced in OPTN-deficient cells

A small fraction of TFRC is constitutively degraded by basal autophagy in many types of cells [21]. But how and why this autophagic degradation of TFRC occurs is not known. It is likely that along with the endosomal membrane, TFRC is also delivered to autophagosomes. Since TFRC is constitutively degraded by autophagy, and OPTN forms a complex with TFRC [15–17], we examined the role of OPTN in TFRC degradation by autophagy. Autophagy was induced by amino acid starvation by incubating the cells in Earle’s balanced salt solution (EBSS). Upon induction of autophagy by amino acid starvation, *Optn^+/+^* cells showed an increase in TFRC level initially and then a decrease after 4h **(Fig. 3A)**. However, this starvation-induced decrease in TFRC level was not seen in *Optn^−/−^* cells **(Fig. 3A)**. Quantification of western blots showed that the level of TFRC decreased significantly after 6 hours of starvation in *Optn^+/+^* cells but not in *Optn^−/−^* cells **(Fig. 3B).** The level of TFRC mRNA was comparable in *Optn^+/+^* and *Optn^−/−^* cells, and upon starvation for 6 hours there was significant increase in TFRC mRNA level in *Optn^+/+^* cells but not in *Optn^−/−^* cells **(Fig. 3C)**. These results suggest that starvation-induced reduction in TFRC protein level seen in *Optn^+/+^* cells was not due to a reduction in mRNA level.

**Figure 3.**
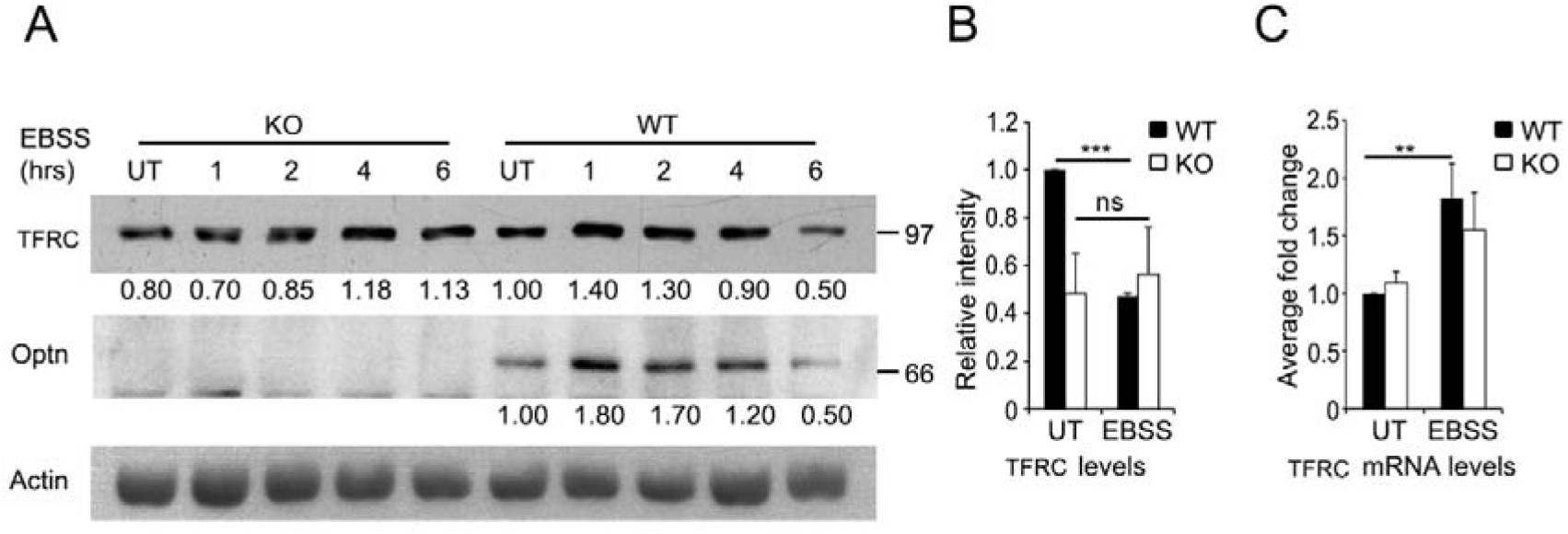
Autophagy mediated TFRC degradation is reduced in optineurin-deficient cells. **(A)** Western blot showing TFRC levels in *Optn^−/−^* (KO) and *Optn^+/+^* (WT) MEFs, upon EBSS treatment for 1-6 hours. Actin was used as loading control. Numbers below the blots indicate relative expression levels after normalization with actin. UT, untreated. **(B)** Bar diagram shows average levels (±SEM) of TFRC protein in *Optn^+/+^* (WT) and *Optn^−/−^* (KO) MEFs, untreated or treated with EBSS for 6 hrs. n=4, ***p<0.001 compared to WT untreated. **(C)** Bar diagram showing average (±SEM) of TFRC mRNA level in WT and KO MEFs, untreated or treated with EBSS for 6 hrs. n=5 for untreated and n=3 for EBSS treated; **p<0.01 compared to WT untreated.

To test the role of OPTN in autophagy mediated degradation of TFRC in a different system we used RGC5, a retinal cell line. RGC5 was used because it has higher level of TFRC as compared to MEFs (Fig. 4A) and it has been shown previously that TFRC is degraded during basal and starvation induced autophagy in this cell line [21]. RGC5 is a retinal ganglion precursor-like cell line [31]. To dissect the role of autophagy in TFRC degradation we knocked down Atg5, a protein required for autophagy, in RGC5 cell line and compared TFRC levels (Fig. 4B). TFRC levels were increased upon ShRNA mediated Atg5 knockdown as compared to control ShRNA transfected sample, providing further evidence for the role of autophagy in TFRC degradation (Fig. 4B). OPTN knockdown in these cells resulted in enhanced TFRC level, and TFRC degradation upon amino acid starvation induced autophagy was reduced in OPTN knockdown cells (Fig. 4C). Knockdown of OPTN also resulted in significantly decreased percentage of TFRC-positive autophagosomes in these cells under basal as well as amino acid starvation induced autophagic condition (Fig. 4D and E). Overall these results suggest a vital role of OPTN in the formation of TFRC-positive autophagosomes, possibly by mediating the delivery of membrane from the TFRC-positive endosomes or recycling endosomes to autophagosomes.

**Figure 4.**
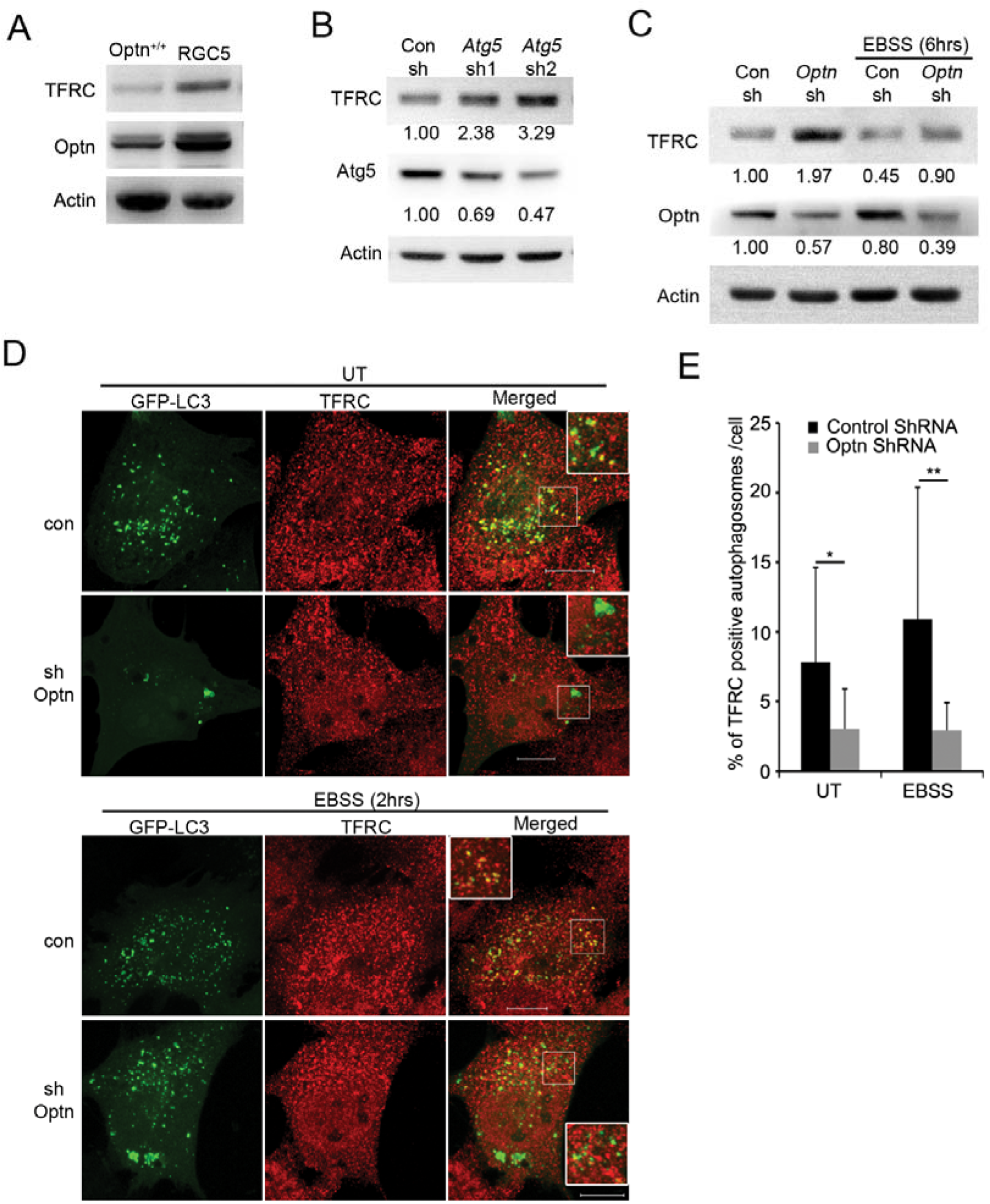
Optineurin mediates formation of TFRC-positive autophagosomes in retinal cells: **(A)** Western blot showing TFRC levels in *Optn^+/+^* MEFs (*Optn^+/+^*) and RGC5 cells. **(B)** Western blot showing increase in TFRC levels upon Atg5 knockdown mediated by two different ShRNAs (*Atg5* sh1 and *Atg5* sh2) and a control shRNA (Con sh) in RGC5 cells. Actin was used as loading control. Numbers below the blots indicate relative expression levels after normalization with actin. **(C)** Western blot showing change in TFRC levels upon ShRNA mediated Optn knockdown in untreated or EBSS treated RGC5 cells. Numbers below the blots indicate relative expression levels after normalization with actin. **(D)** Representative confocal images showing TFRC-positive autophagosomes (GFP-LC3 positive dots) in RGC5 cells transfected with GFP-LC3B along with control ShRNA or *Optn* ShRNA, untreated or treated with EBSS for 2 hrs. Magnified area is shown as inset. Scale bar: 10μm. **(E)** Bar diagram representing percentage of TFRC-positive autophagosomes in RGC5 cells either transfected with control ShRNA or *Optn* ShRNA, untreated or treated with EBSS for 2 hrs. n=10 cells. *p<0.05, **p<0.01.

### A disease-associated mutant E478G is defective in the formation of TFRC-positive autophagosomes

Mutations in OPTN are associated with normal tension glaucoma and ALS (Amyotrophic lateral sclerosis) [32, 33]. ALS-associated mutations of OPTN include truncation, deletion and missense mutations, and it has been suggested that loss of function as well as gain of function mechanisms are likely to be involved in ALS pathogenesis caused by OPTN mutations. One of the mutations identified in the original study was E478G, which was suggested to be a disease causing mutation [33, 34]. Unlike wild type OPTN, this mutant was unable to promote maturation of phagophore into autophagosome. This mutation is located in ubiquitin binding domain (UBD) of OPTN and it abolished binding of OPTN to ubiquitin [7, 30]. We examined its role in promoting formation of TFRC-positive autophagosomes. *Optn^−/−^* MEFs were transfected with GFP-LC3B along with HA-tagged wild type OPTN (WT-OPTN) or E478G mutant or control plasmid. The fixed cells were stained with HA and TFRC antibodies. Unlike WT-OPTN, this mutant was unable to increase the percentage of TFRC positive autophagosomes when overexpressed in *Optn^−/−^* MEFs (Fig. 5E and F). These results suggest that the E478G mutant is defective in the delivery of membrane from TFRC-positive endosomes to autophagosomes, thus ubiquitin binding capacity of OPTN is prerequisite for the formation of TFRC-positive autophagosomes.

**Figure 5.**
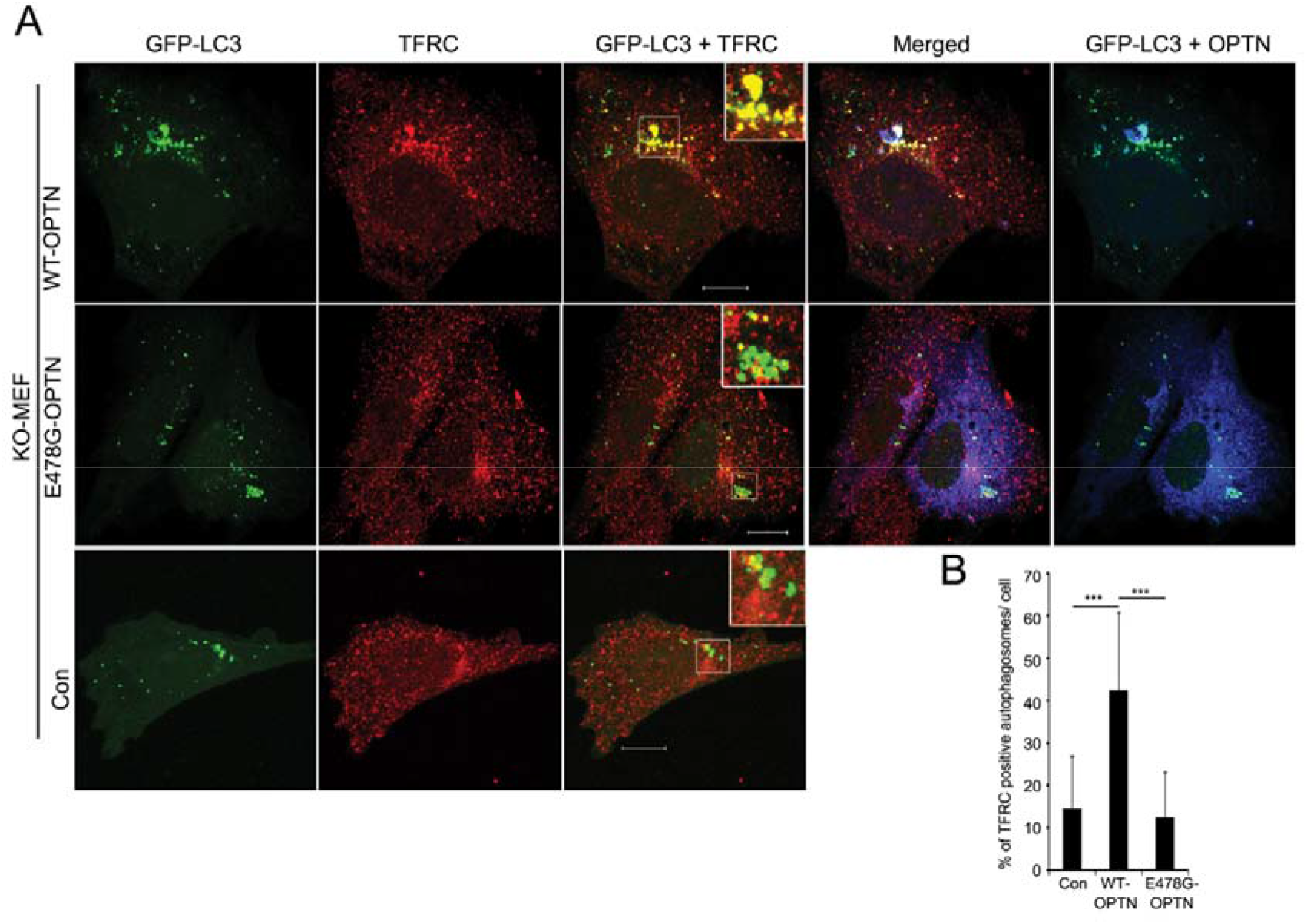
ALS-associated mutant E478G is defective in the formation of TFRC-positive autophagosomes. **(A)** Representative confocal images showing TFRC-positive autophagosomes in *Optn^−/−^* (KO) MEFs transfected with GFP-LC3B along with either control plasmid (Con) or HA-WT-OPTN or HA-E478G-OPTN mutant. TFRC was stained with anti-mouse Cy3 antibody (represented as red) while HA-OPTN or HA-E478G was stained with HA antibody followed by anti-rabbit Alexa-633 antibody (represented as blue). Magnified area is shown as inset. Scale bar: 10μm. **(B)** Bar diagram showing percentage of TFRC-positive autophagosomes per cell in *Optn^−/−^* MEFs transfected with GFP-LC3B along with control plasmid (Con) or WT-OPTN or E478G-OPTN. n=18 cells. ***p<0.001.

## Discussion

Various membranous organelles, including ER, plasma membrane, Golgi and TFRC-positive recycling endosomes contribute membrane for the biogenesis of autophagosomes [27, 28, 35–39]. How the membrane is delivered to autophagosome, is not clear. Here, we investigated the role of OPTN in the delivery of membrane from TFRC-positive endosomes to autophagosomes. Our results suggest that OPTN mediates delivery of TFRC (and possibly associated membrane) from TFRC-positive endosomes or recycling endosomes to autophagosomes. This conclusion is supported by our results showing that (a) the percentage of TFRC-positive autophagosomes is reduced in OPTN deficient cells, and overexpressed OPTN increases the formation of TFRC-positive autophagosomes; and (b) TFRC degradation by autophagy is dependent on OPTN. Reduced degradation of TFRC in OPTN-deficient cells is likely to be due to reduced delivery of TFRC to autophagosomes. Since vesicle trafficking is a multi-step process, the requirement of OPTN for delivery of TFRC to autophagosomes could be for any one of the steps involved in this delivery process.

Previously it has been shown that OPTN, an effector of Rab8, is involved in mediating endocytic trafficking of TFRC-positive endosomes from the periphery of the cell to the juxtanuclear region. Overexpressed as well as endogenous OPTN forms a complex with TFRC [15–17]. It seems likely that trafficking of TFRC and associated membrane from endosomes or recycling endosomes to autophagosomes is facilitated by formation of a complex of optineurin with TFRC.

OPTN performs various autophagic functions in the cell. It is an autophagy receptor that mediates cargo-selective autophagy of bacteria, damaged mitochondria and mutant protein aggregates [7, 9, 10]. It is also involved in non-selective (basal and starvation-induced) autophagy, where it promotes maturation of phagophore into autophagosome. In OPTN-deficient cells, phagophores accumulate whereas production of autophagosomal protein LC3-II (the lapidated form of LC3) from LC3-I is reduced [30]. OPTN promotes autophagosome formation by increasing production of LC3-II from LC3-I by enhancing recruitment of Atg-12-5-16L1 complex to the phagophore [30]. Here, we suggest that the delivery of membrane from TFRC-positive endosomes to autophagosomes by OPTN may also contribute to autophagosome formation. This is shown schematically in Fig. 6.

**Figure 6.**
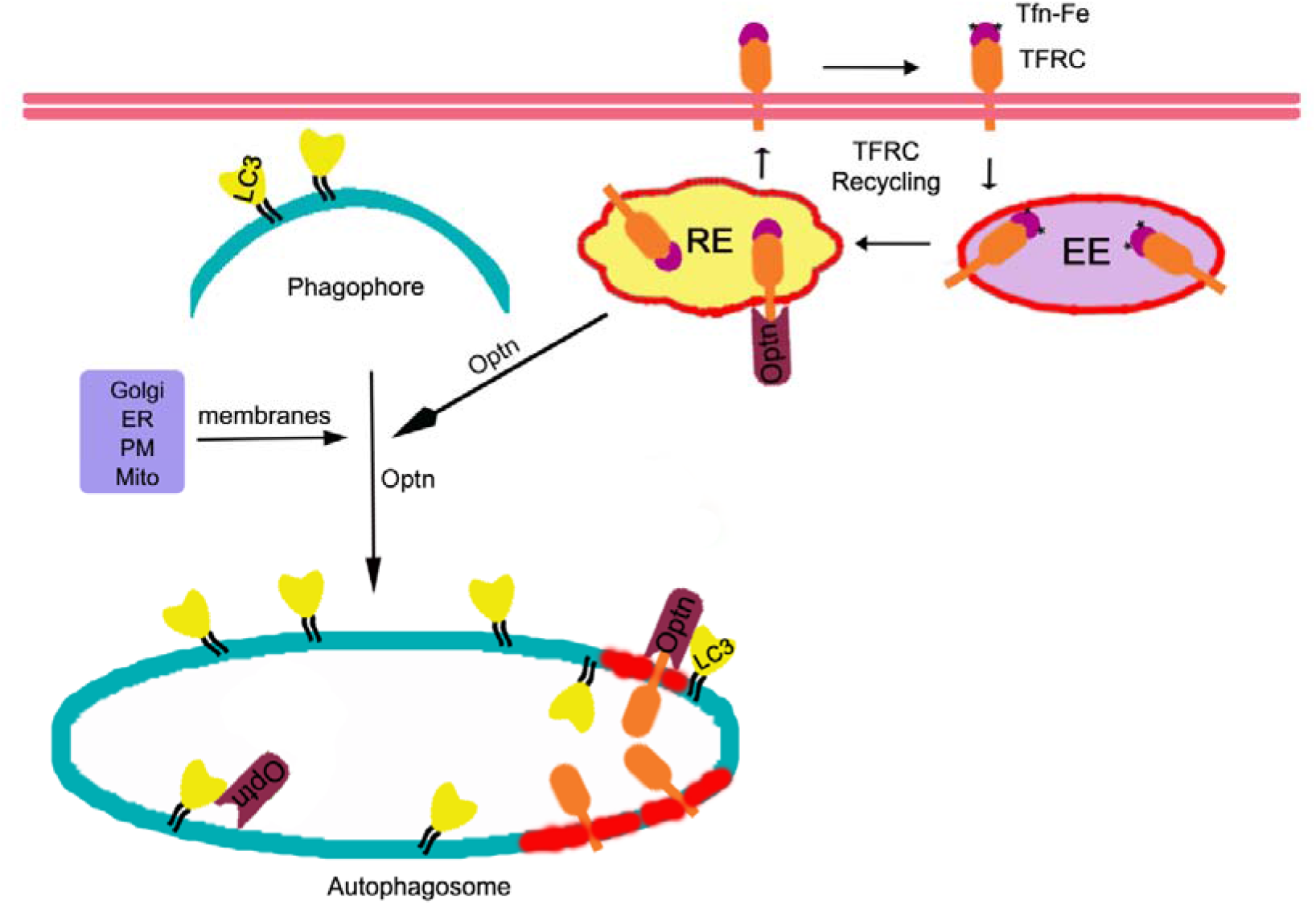
Model depicting role of optineurin in membrane delivery to autophagosome. During autophagosome formation membrane is contributed by various organelles like ER, mitochondria, Golgi, plasma membrane and recycling endosome (RE). Here, we suggest that OPTN, which is known to play an important role in endocytic trafficking and recycling of TFRC, mediates delivery of TFRC along with membrane from recycling endosomes to forming autophagosomes. OPTN is known to promote maturation of phagophore into autophagosome by enhancing recruitment of Atg-12-5-16L1 complex to phagophore leading to increased LC3-II production [30]. Thus, we suggest that OPTN contributes to autophagosome formation through these two processes, enhancement of LC3-II production (reported earlier by us [30]) and membrane delivery from TFRC-positive endosomes. EE, early endosome; Fe, iron; RE, recycling endosome; Tfn, transferrin; PM, plasma membrane; Ub, ubiquitin.

Mutations of OPTN are associated with two neurodegenerative diseases, adult onset primary open angle glaucoma and ALS. ALS-associated mutants of OPTN include deletion, truncation and missense mutations. Loss of function as well as gain of function mechanisms have been proposed to explain pathogenesis due to OPTN mutations. E478G is an ALS-associated mutation defective in ubiquitin binding that is also defective in autophagosome formation. Our results show that this mutant is defective in promoting formation of TFRC-positive autophagosomes. This suggests that ubiquitin-binding function of OPTN is required for delivery of membrane from TFRC-positive endosomes to autophagosome.

## Materials and methods

### Cell culture and transfection

*Optn^+/+^* (WT) and *Optn^−/−^* (KO) MEFs were generated in our laboratory as described [30, 40]. MEFs and RGC5 cells were grown at 37°C in DMEM supplemented with 10% fetal bovine serum in a humidified incubator with 5% CO2. For MEFs, DMEM was supplemented with 0.1mM β-mercaptoethanol, 50 IU/ml penicillin and 50μg/ml streptomycin. Transfections were performed using Lipofectamine (Invitrogen, 18324-012) and Lipofectamine PLUSTM (Invitrogen, 11514-015) reagents, or Lipofectamine 2000 (invitrogen, 11669-019) reagent by following manufacturer’s instructions. For inducing starvation, cells were washed three times with PBS, and then incubated in Earle’s balanced salt solution (EBSS, Invitrogen, 14155063) containing CaCl_2_ (1.8mM) and MgSO_4_ (0.81 mM) at 37°C for indicated time.

### Expression Vectors

Plasmids for expressing HA tagged human OPTN and E478G mutant of OPTN have been described earlier [30, 41]. Plasmids for expressing mouse Atg5 shRNA and Optn ShRNA have been described [21, 42]. GFP-LC3B construct was kindly provided by Dr. Terje Johansen (University of Tromsø, Tromsø, Norway) [43].

### Antibodies and reagents

Rabbit polyclonal antibodies-anti-optineurin (Abcam, ab23666), anti-LC3B (CST, 27755), anti-HA (Santa Cruz, sc-805) and anti-ATG5 (CST, 8540) were used. Mouse monoclonal antibodies-anti-TFRC (Invitrogen, 136800) and anti-actin (Millipore, MAB1501) were used.

### Immunofluorescence microscopy

Cells grown on glass coverslips were transfected with required plasmids. After indicated time, cells were fixed in 3.7% formaldehyde for 10 min and then permeabilised using 0.5% Triton X-100 and 0.05% Tween-20 for 6 min. Immunofluorescence staining was carried out essentially as described [30]. For immunostaining of endogenous LC3, cells were fixed with 3% paraformaldehyde for 20 min, followed by aldehyde quenching with 50mM NH_4_Cl for 10 min (39). Mounting medium with DAPI was used for visualising the nucleus. The images were acquired using LSM510 meta NLO confocal microscope from Carl Ziess (Jena Germany) or Leica TCS SP8 with 63X/1.4NA oil immersion objective lens. Two middle optical Z-sections of 0.35μm thickness were used for projection.

### Quantification of puncta

Number and size of puncta were quantified using ImageJ (NIH) software. Optical Z-sections of 0.35μm thickness of the cell were taken using confocal microscope with 63X oil immersion objective lens. For the quantification of autophagy puncta 8-10 optical Z-sections were orthogonally projected to obtain a composite image, and puncta with area of 0.1-3.0 cm^2^ were quantitated. To calculate double positive puncta, those puncta that overlapped by at least 0.1cm^2^ of area were counted.

### Quantitative PCR

Total RNA was isolated from MEFs using Trizol reagent (Invitrogen). cDNA synthesis was carried out by using SuperScript III First-Strand Synthesis System (Invitrogen) after DNase I (NEB) treatment of RNA. Quantitative PCR was performed by using 7900HT fast real time PCR system (Applied Biosystems). Expression of TFRC gene was normalized with actin.

### Statistical analysis

Bar diagrams represent average ± standard error of mean (SEM) or standard deviation (SD). Statistical significance between averages was calculated using Student’s T-test (two tailed). P value less than 0.05 was considered significant.

## Author contributions

MB and KS planned, performed and analysed most of the experiments and helped in writing the manuscript; SM helped in some experiments; GS planned experiments, wrote the manuscript and obtained funding.

## Acknowledgements

GS gratefully acknowledges Indian National Science Academy for Senior Scientist position, and Department of Science and Technology, Government of India for J.C. Bose National Fellowship (grant no: SR/S2/JCB-41/2010).

## References

1. Ying, H. & Yue, B. Y. Cellular and molecular biology of optineurin, Int Rev Cell Mol Biol. 2012;294, 223–258.

2. Kachaner, D., Genin, P., Laplantine, E. & Weil, R. Toward an integrative view of Optineurin functions in Cell Cycle 2012;11(15):2808–2818.

3. Minegishi, Y., Nakayama, M., Iejima, D., Kawase, K. & Iwata, T. (2016) Significance of optineurin mutations in glaucoma and other diseases, Progress in retinal and eye research. 2016;55, 149–181.

4. Markovinovic, A., Cimbro, R., Ljutic, T., Kriz, J., Rogelj, B. & Munitic, I. Optineurin in amyotrophic lateral sclerosis: Multifunctional adaptor protein at the crossroads of different neuroprotective mechanisms, Prog Neurobiol. 2017;154, 1–20.

5. Toth, R. P. & Atkin, J. D. Dysfunction of Optineurin in Amyotrophic Lateral Sclerosis and Glaucoma, Front Immunol. 2018; 9, 1017.

6. Swarup, G. & Sayyad, Z. Altered Functions and Interactions of Glaucoma-Associated Mutants of Optineurin, Front Immunol. 2018; 9, 1287.

7. Wild, P., Farhan, H., McEwan, D. G., Wagner, S., Rogov, V. V., Brady, N. R., Richter, B., Korac, J., Waidmann, O., Choudhary, C., Dotsch, V., Bumann, D. & Dikic, I. Phosphorylation of the autophagy receptor optineurin restricts Salmonella growth, Science. 2011; 333, 228–233.

8. Ramachandran G, Moharir SC, Raghunand TR, Swarup G. Biochem Biophys Res Commun. Optineurin modulates ER stress-induced signaling pathways and cell death. 2021;534: 297–302.

9. Korac, J., Schaeffer, V., Kovacevic, I., Clement, A. M., Jungblut, B., Behl, C., Terzic, J. & Dikic, I. Ubiquitin-independent function of optineurin in autophagic clearance of protein aggregates, J Cell Sci. 2013; 126, 580–592.

10. Wong, Y. C. & Holzbaur, E. L. Optineurin is an autophagy receptor for damaged mitochondria in parkin-mediated mitophagy that is disrupted by an ALS-linked mutation, Proceedings of the National Academy of Sciences of the United States of America. 2014; 111: E4439–4448.

11. Shen WC, Li HY, Chen GC, Chern Y, Tu PH. Mutations in the ubiquitin-binding domain of OPTN/optineurin interfere with autophagy-mediated degradation of misfolded proteins by a dominant-negative mechanism. Autophagy. 2015;11(4):685–700.

12. Hattula, K, and Peranen, J. FIP-2, a coiled-coil protein, links Huntingtin to Rab8 and modulates cellular morphogenesis. Curr Biol 2000; 10:1603–1606.

13. Hattula K, Furuhjelm J, Tikkanen J, Tanhuanpää K, Laakkonen P, Peränen J. Characterization of the Rab8-specific membrane traffic route linked to protrusion formation. J Cell Sci 2006; 119:4866–4877.

14. Sahlender DA, Roberts RC, Arden SD, Spudich G, Taylor MJ, Luzio JP, Kendrick-Jones J, Buss F. Optineurin links myosin VI to the Golgi complex and is involved in Golgi organization and exocytosis. J Cell Biol 2005;169:285–295.

15. Nagabhushana A, Chalasani ML, Jain N, Radha V, Rangaraj N, Balasubramanian D, Swarup G. Regulation of endocytic trafficking of transferrin receptor by optineurin and its impairment by a glaucoma-associated mutant. BMC Cell Biol. 2010;11:4.

16. Park B, Ying H, Shen X, Park JS, Qiu Y, Shyam R, Yue BY. Impairment of protein trafficking upon overexpression and mutation of optineurin. PLoS One 2010;5(7):e11547.

17. Vaibhava V, Nagabhushana A, Chalasani ML, Sudhakar C, Kumari A, Swarup G. Optineurin mediates a negative regulation of Rab8 by the GTPase-activating protein TBC1D17. J Cell Sci 2012;125:5026–5039.

18. Optineurin: A Coordinator of Membrane-Associated Cargo Trafficking and Autophagy. Ryan TA, Tumbarello DA. Front Immunol. 2018;9: 1024.

19. MacKenzie EL, Iwasaki K, Tsuji Y. Intracellular iron transport and storage: from molecular mechanisms to health implications. Antioxid Redox Signal 2008;10:997–1030.

20. Lee DA, Goodfellow JM. The pH-induced release of iron from transferrin investigated with a continuum electrostatic model. Biophys J 1998; 74:2747–59. doi:10.1016/S0006-3495(98)77983-4.

21. Sirohi K, Chalasani ML, Sudhakar C, Kumari A, Radha V, Swarup G. M98K-OPTN induces transferrin receptor degradation and RAB12-mediated autophagic death in retinal ganglion cells. Autophagy 2013; 9(4):510–527.

22. Matsui T, Itoh T, Fukuda M. Small GTPase Rab12 regulates constitutive degradation of transferrin receptor. Traffic 2011; 12:1432–1443.

23. Yang Z & Klionsky DJ (2009) An overview of the molecular mechanism of autophagy. Curr. Top. Microbiol. Immunol. 335:1–32.

24. Stolz A, Ernst A, & Dikic I (2014) Cargo recognition and trafficking in selective autophagy. Nat. Cell Biol. 16(6):495–501.

25. Levine B, Packer M, & Codogno P (2015) Development of autophagy inducers in clinical medicine. J. Clin. Invest. 125(1):14–24.

26. Tsukada M & Ohsumi Y (1993) Isolation and characterization of autophagy-defective mutants of Saccharomyces cerevisiae. FEBS Lett. 333(1-2):169–174.

27. Mizushima N (2010) The role of the Atg1/ULK1 complex in autophagy regulation. Curr. Opin. Cell Biol. 22(2):132–139.

28. Tooze SA, Abada A, Elazar Z. Endocytosis and autophagy: exploitation or cooperation? Cold Spring Harb Perspect Biol 2014; 6:a018358.

29. Longatti A, Lamb CA, Razi M, Yoshimura S, Barr FA, Tooze SA. TBC1D14 regulates autophagosome formation via Rab11- and ULK1-positive recycling endosomes. J Cell Biol 2012; 197:659–675.

30. Bansal, M., Moharir, S. C., Sailasree, S. P., Sirohi, K., Sudhakar, C., Sarathi, D. P., Lakshmi, B. J., Buono, M., Kumar, S. & Swarup, G. (2018) Optineurin promotes autophagosome formation by recruiting the autophagy-related Atg12-5-16L1 complex to phagophores containing the Wipi2 protein, J Biol Chem. 293, 132–147.

31. Sayyad, Z., Sirohi, K., Radha, V., Swarup, G. (2017) 661W is a retinal ganglion precursor-like cell line in which glaucoma-associated optineurin mutants induce cell death selectively. Sci Rep 7, 16855.

32. Rezaie, T., Child, A., Hitchings, R., Brice, G., Miller, L., Coca-Prados, M., Heon, E., Krupin, T., Ritch, R., Kreutzer, D., Crick, R. P. & Sarfarazi, M. (2002) Adult-onset primary open-angle glaucoma caused by mutations in optineurin, Science. 295, 1077–1079.

33. Maruyama, H., Morino, H., Ito, H., Izumi, Y., Kato, H., Watanabe, Y., Kinoshita, Y., Kamada, M., Nodera, H., Suzuki, H., Komure, O., Matsuura, S., Kobatake, K., Morimoto, N., Abe, K., Suzuki, N., Aoki, M., Kawata, A., Hirai, T., Kato, T., Ogasawara, K., Hirano, A., Takumi, T., Kusaka, H., Hagiwara, K., Kaji, R. & Kawakami, H. (2010) Mutations of optineurin in amyotrophic lateral sclerosis, Nature. 465, 223–226.

34. Ito H, Nakamura M, Komure O, Ayaki T, Wate R, Maruyama H, Nakamura Y, Fujita K, Kaneko S, Okamoto Y, et al. Clinicopathologic study on an ALS family with a heterozygous E478G optineurin mutation. Acta Neuropathol 2011; 122:223–229.

35. Razi M, Chan EY, Tooze SA. Early endosomes and endosomal coatomer are required for autophagy. J Cell Biol 2009; 185:305–321.

36. Ravikumar B, Moreau K, Jahreiss L, Puri C, Rubinsztein DC. Plasma membrane contributes to the formation of pre-autophagosomal structures. Nat Cell Biol 2010; 12:747–757.

37. Hailey DW, Rambold AS, Satpute-Krishnan P, Mitra K, Sougrat R, Kim PK, Lippincott-Schwartz J. Mitochondria supply membranes for autophagosome biogenesis during starvation. Cell 2010; 141:656–667.

38. Axe EL, Walker SA, Manifava M, Chandra P, Roderick HL, Habermann A, Griffiths G, Ktistakis NT. Autophagosome formation from membrane compartments enriched in phosphatidylinositol 3-phosphate and dynamically connected to the endoplasmic reticulum. J Cell Biol 2008; 182:685–701.

39. Geng J, Nair U, Yasumura-Yorimitsu K, Klionsky DJ. Post-Golgi Sec proteins are required for autophagy in Saccharomyces cerevisiae. Mol Biol Cell 2010; 21:2257–2269.

40. Moharir SC, Bansal M, Ramachandran G, Ramaswamy R, Rawat S, Raychaudhuri S, Swarup G. Identification of a splice variant of optineurin which is defective in autophagy and phosphorylation. Biochim. Biophys. Acta Mol. Cell Res. 2018;1865:1526–1538,

41. Chalasani ML, Radha V, Gupta V, Agarwal N, Balasubramanian D, Swarup G. A glaucoma-associated mutant of optineurin selectively induces death of retinal ganglion cells which is inhibited by antioxidants. Invest. Ophthalmol. Vis. Sci. 2007; 48(4):1607–1614.

42. Sirohi K, Kumari A, Radha V & Swarup G (2015) A glaucoma-associated variant of optineurin, M98K, activates Tbk1 to enhance autophagosome formation and retinal cell death dependent on Ser177 phosphorylation of optineurin. PLoS One 2015;10, e0138289.

43. Pankiv S, Clausen TH, Lamark T, Brech A, Bruun JA, Outzen H, Øvervatn A, Bjørkøy G, Johansen T. p62/SQSTM1 binds directly to Atg8/LC3 to facilitate degradation of ubiquitinated protein aggregates by autophagy. J. Biol. Chem. 2007; 282(33):24131–24145.

